# Maturation of the human B-cell receptor repertoire with age

**DOI:** 10.1101/609651

**Authors:** Marie Ghraichy, Jacob D. Galson, Aleksandr Kovaltsuk, Valentin von Niederhäusern, Jana Pachlopnik Schmid, Mike Recher, Annaïse J Jauch, Enkelejda Miho, Dominic F. Kelly, Charlotte M. Deane, Johannes Trück

## Abstract

B cells play a central role in adaptive immune processes, mainly through the production of antibodies. The maturation of the B-cell system with age is poorly studied. We extensively investigated age-related alterations of naïve and antigen-experienced B-cell receptor (BCR) repertoires. The most significant changes were observed in the first 10 years of life, and were characterized by altered immunoglobulin gene usage and an increased frequency of mutated antibodies structurally diverging from their germline precursors. Older age was associated with an increased usage of downstream constant region genes and fewer antibodies with self-reactive properties. As mutations accumulated with age, the frequency of germline-encoded self-reactive antibodies decreased, indicating a possible beneficial role of self-reactive B-cells in the developing immune system. Our results suggest a continuous process of change through childhood across a broad range of parameters characterizing BCR repertoires and stress the importance of using well-selected, age-appropriate controls in BCR studies.

## Introduction

B cells play a central role in physiological adaptive immune processes and exert their main effector function through production of antibodies (1). B cells also contribute to the pathogenesis of autoimmune disease via generation of auto-reactive antibodies and modulation of T-cell responses (2, 3). The heavy and light chains of the B-cell receptor (BCR, membrane-bound antibody) are generated in the bone marrow by recombining individual variable (V), diversity (D) and joining (J) gene segments through a process called VDJ recombination. Upon antigen recognition, a BCR is further diversified through rounds of somatic hypermutation (SHM) leading to affinity maturation whereby B cells with improved antigen-binding properties are selected in the germinal center. Class switch recombination (CSR) is also initiated following antigen encounter, causing a change in the constant region of the BCR and in its effector function.

Detailed characterization of B cells and their respective BCR sequences offers important information on B-cell generation and selection as well as immune competence in health and disease. High-throughput sequencing of antibody genes (Ig-seq) has become a widely used tool in human translational research (4, 5). Abnormal B-cell responses can be explored by investigating BCR repertoires from patients and comparing their characteristics to those of healthy controls. The limited data already available suggest that significant changes occur in the properties of BCR repertoires with age (6). It is therefore important to establish robust data on normal BCR repertoires within sufficiently narrow age-bands to fully understand the process of BCR maturation. This will facilitate the use of Ig-seq to understand changes of relevance to childhood disease. Given the high burden of infectious diseases in childhood and the importance of effective immune response to vaccines to prevent infection, this is an important group from which to have normative data. There are very few studies that have used Ig-seq to investigate the healthy BCR repertoire, and these studies include a limited age range of participants (7–10). In a more detailed study, Ijspeert et al. reported on the antigen-experienced (i.e. IgA and IgG) BCR repertoires of 38 healthy control (HC) samples with their ages ranging from newborn to 74 years (11). The authors found several characteristics of the studied BCR repertoire varying with age and identified patterns that are specific for isotype subclasses. However, their study was limited by the number of samples from children, the low depth of sequencing, and the small number of B-cell subsets analyzed.

We aimed to assess in detail the naïve and antigen-experienced BCR repertoires in children and young adults using isotype-resolved barcoded RNA-based Ig-seq technology and extensive bioinformatic analysis. This approach allowed us to comprehensively address the age effect on the BCR repertoire in healthy individuals and also provides a robust data set that can serve as a future reference for studying BCR repertoires in children as well as young adults with disease.

## Methods

### Study participants and cell isolation

Study participants were recruited with informed consent under ethical approval (KEK-ZH 2015-0555 and EKNZ 2015-187). Blood samples (5-9 mL) were collected at a single time point from 53 healthy participants aged 6 months to 50 years (**Supplementary table 1**). Peripheral blood mononuclear cells (PBMC) were isolated by centrifugation of PBS-diluted blood over Ficoll-Paque Plus (Sigma-Aldrich). Either PBMC or B cells magnetically sorted using the AutoMACS Pro cell separator and CD19+ microbeads (both Miltenyi Biotec), were lysed in RLT buffer (Qiagen), snap frozen on dry ice and then stored at -80 °C prior to use. Cells were counted using an optical microscope and an improved Neubauer chamber. The B-cell number was recorded based on actual counts or estimated using PBMC counts and either B-cell frequencies from flow cytometry performed on the same blood sample or the median percentage of age-dependent reference values (12) if the former was not available.

### RNA isolation and library preparation

RNA was extracted from stored samples using the RNeasy Mini Kit (Qiagen). Reverse transcription was performed using SuperScript III/IV (Invitrogen) according to the manufacturer’s instructions and constant region primers that included 14 nt unique molecular identifiers (UMI), and partial p7 adaptors. Two reverse transcription reactions were carried out for each sample: one with a mix of IgM and IgD-specific reverse primers and another with a mix of IgA, IgG, and IgE-specific reverse primers. From 6 samples, one mix with all C region primers were used in a single reaction. Primer sequences with concentrations are included in **Supplementary table 2**. BCR heavy chain rearrangements were amplified in a two-round multiplex PCR; the first round using a mix of V family specific forward primers with partial p5 adaptors, and the second round to complete the adaptor sequences. PCR conditions for the first round were 95 °C for 5 min, either 8 cycles (IgD/IgM) or 12 cycles (IgA/E/G) of 98 °C for 20s, 60 °C for 45s and 72 °C for 1 min, and 72 °C for 5 minutes. The PCR conditions for the second round were 95 °C for 5 min, 22 cycles of 98 °C for 20s, 69 °C for 20s and 72 °C for 15 sec, and 72 °C for 5 minutes. PCR amplicons were gel-extracted, purified and quantified using the Illumina qPCR library quantification protocol. Individual libraries were normalized based on concentration and then multiplexed in batches of 24 for sequencing on the Illumina MiSeq platform (2 × 300 bp paired-end chemistry). Compared with short-read sequencing protocols, long read RNA sequencing on the Illumina MiSeq instrument results in less deep sequencing. However, long read sequencing provides information on the entire VDJ sequence and the constant region of the BCR allowing for accurate distinction between isotypes and subclasses.

### Sequence processing, annotation and somatic hypermutation

Samples were demultiplexed via their Illumina indices, and initially processed using the Immcantation toolkit (13, 14). Briefly, raw fastq files were filtered based on a quality score threshold of 20. Paired reads were joined if they had a minimum length of 10 nt, maximum error rate of 0.3 and a significance threshold of 0.0001. Reads with identical UMI (i.e. originating from the same mRNA molecule) were collapsed to a consensus sequence. Reads with identical full-length sequence and identical constant primer but differing UMI were further collapsed resulting in a dataset containing a set of unique sequences per sample and isotype. Sequences were then submitted to IgBlast (15) for VDJ assignment and sequence annotation, and unproductive sequences removed. Constant region sequences were mapped to germline using Stampy (16) for isotype (subclass) annotation, and only sequences with a defined constant region were kept for further analysis. The number and type of V gene mutations was calculated using the shazam R package (14). Levels of somatic hypermutation (SHM) were determined by calculating V gene mutations in individual sequences, and mean values were calculated across samples and cell subsets.

### Sequence clustering, clonal lineages and antigen-driven selection

Sequences were independently clustered for each sample to group together those arising from clonally related B cells. The clustering required identical V and J segment use, identical complementary-determining region (CDR) 3 length, and allowing a 1 in 15 nucleotides mismatch in the CDR3 as previously determined (7). Lineages were constructed from clusters using the alakazam R package (17). To account for read depth variation, lineage trees were constructed on subsamples of the original data. Specifically, we randomly sampled 25’609 sequences (corresponding to the lowest number of reads available for a sample) from every HC sample. For calculation of selection pressure of samples, individual sequences within clusters are not independent events, so an effective representative sequence of each clonal group was determined using the default settings of shazam. Selection pressure was calculated using BASELINe (18) implemented within shazam. The statistical framework used to test for selection was CDR_R / (CDR_R + CDR_S), which normalizes for the observed increase in the total number of mutations with age. The replacement/silent (R/S) mutation ratio was measured separately in framework regions (FWRs) and CDRs. In sequences with replacement but no silent mutations, the number of silent mutations was set to 1.

### From Sequence to Structure

The SAAB+ pipeline was employed to annotate BCR repertoires with structural information (19). Briefly, BCR repertoires were numbered with the IMGT scheme (20) and filtered for structural viability using ‘ANARCI parsing’ (21) as per the first steps of the ABOSS algorithm (22). Sequences were filtered out that (i) could not be aligned to the human Hidden Markov Model (HMM) profile of an IMGT germline (ii) had a J gene sequence identity of less than 50% to a human IMGT germline or (iii) contained non-amino acid entries in CDRs. Since the primer masking step in pRESTO (13) can remove the first framework region and positions 127 and 128 of some sequences, ANARCI parsing was customized to account for these exceptions. To retain as many sequences as possible for structural annotation, we substituted undetermined residues in the framework region with the residues from their respective parent germline genes.

To annotate the numbered sequences with canonical loop class information, SAAB+ employs SCALOP (23) with the IMGT CDR definition (20). The expected coverage of canonical loop class sequences with SCALOP is 93%, where 89% of predicted templates will have root-mean-square deviation (RMSD) values for the backbone atoms within 1.5 Å of the correct structure. The SCALOP database dated July 2018 was used in this study.

SAAB+ employs FREAD (24) to annotate CDR-H3 loops with the Protein Data Bank (PDB) code (25) of the closest crystallographically-solved CDR-H3 structure (template). Only CDR-H3 sequences with loop lengths between 5 and 16 were investigated. The expected average RMSD of CDR-H3 template prediction for the human BCR repertoire data is 2.8 Å, with an expected coverage of 48% (19). PDB templates within a 0.6 Å RMSD radius were clustered together (19), reducing 2’943 PDB templates to 1’169 CDR-H3 PDB clusters.

### Statistical analysis and graphing

Statistical analysis and plotting were performed using R (26); all plots were produced using the ggplot2 and ggpubr packages (27, 28). Heatmaps were visualized using the ComplexHeatmap R package (29). PCA plots were created using the R package factoextra (30). In cases where a model was fitted to the data, the R squared of the model and the p value of the chi-squared goodness-of-fit test are shown in the bottom right of the graphs. Other specific tests used are detailed in the figure legends.

### Classification of sequences into cell subsets using isotype and number of mutations

Using constant region annotation and mutation number, individual sequences were grouped into biologically different subsets based on known B-cell subpopulations. Based on the frequency distribution of mutations for IgD and IgM sequences, those with up to 2 nt mutations across the entire V gene were considered “unmutated” (naïve) to account for allelic variance (31) and remaining PCR and sequencing bias (**Supplementary figure 1**). All class-switched sequences were defined as antigen-experienced irrespective of their V gene mutation count. Because of very low sequence numbers, IgE and IgG4 transcripts were excluded from most analysis. The number of sequences of the different subsets among total transcripts by individual are found in **Supplementary table 1**.

**Figure 1:**
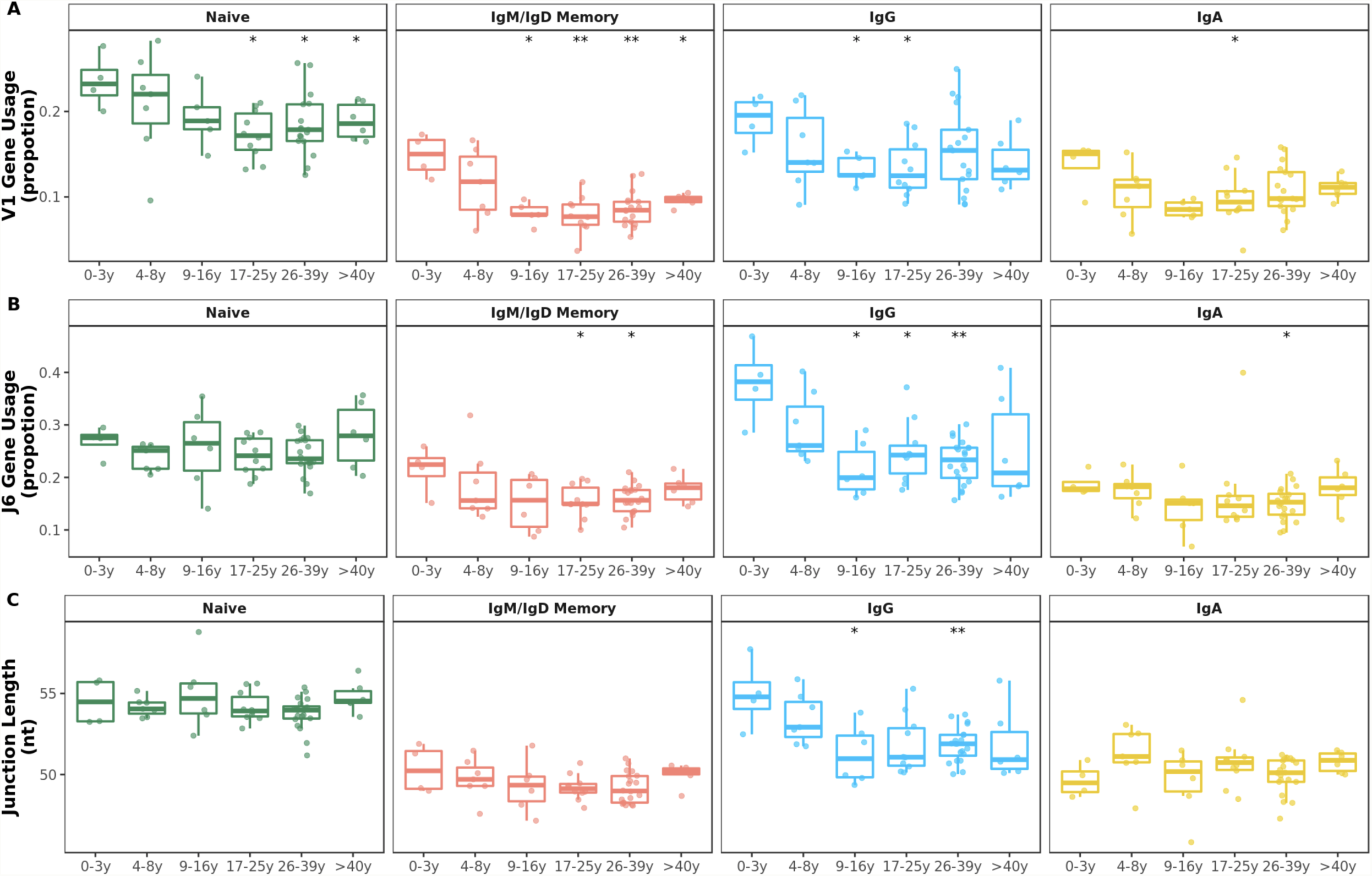
V family and J gene usage changes in early childhood. *A* V1 family usage was significantly reduced in older compared with younger individuals in all BCR repertoires. *B* J6 gene usage significantly decreased during the first 10 years of life mostly in IgG subsets. *C* Mean junction length significantly decreased in the first 10 years of life exclusively in IgG subsets. Comparison of each age group to the 0-3y group was performed using the Wilcoxon test. *p<0.05, **p<0.01

### Data availability

Raw sequence data used for analysis in this study are available at the NCBI Sequencing Read Archive (www.ncbi.nlm.nih.gov/sra) under BioProject number PRJNA527941 including metadata meeting MiAIRR standards (32). The processed and annotated final dataset is available in Zenodo (https://doi.org/10.5281/zenodo.3585046) along with the protocol describing the exact processing steps with the software tools and version numbers.

## Results

We obtained 78’702’939 raw sequences from samples of 53 healthy study participants. Processing, filtering and collapsing resulted in a final dataset of 8’341’669 unique BCR sequences used for downstream analysis. The numbers of unique sequences were significantly reduced after UMI-based collapsing resulting in a correlation with the B-cell numbers per sample (**Supplementary figure 2**).

**Figure 2:**
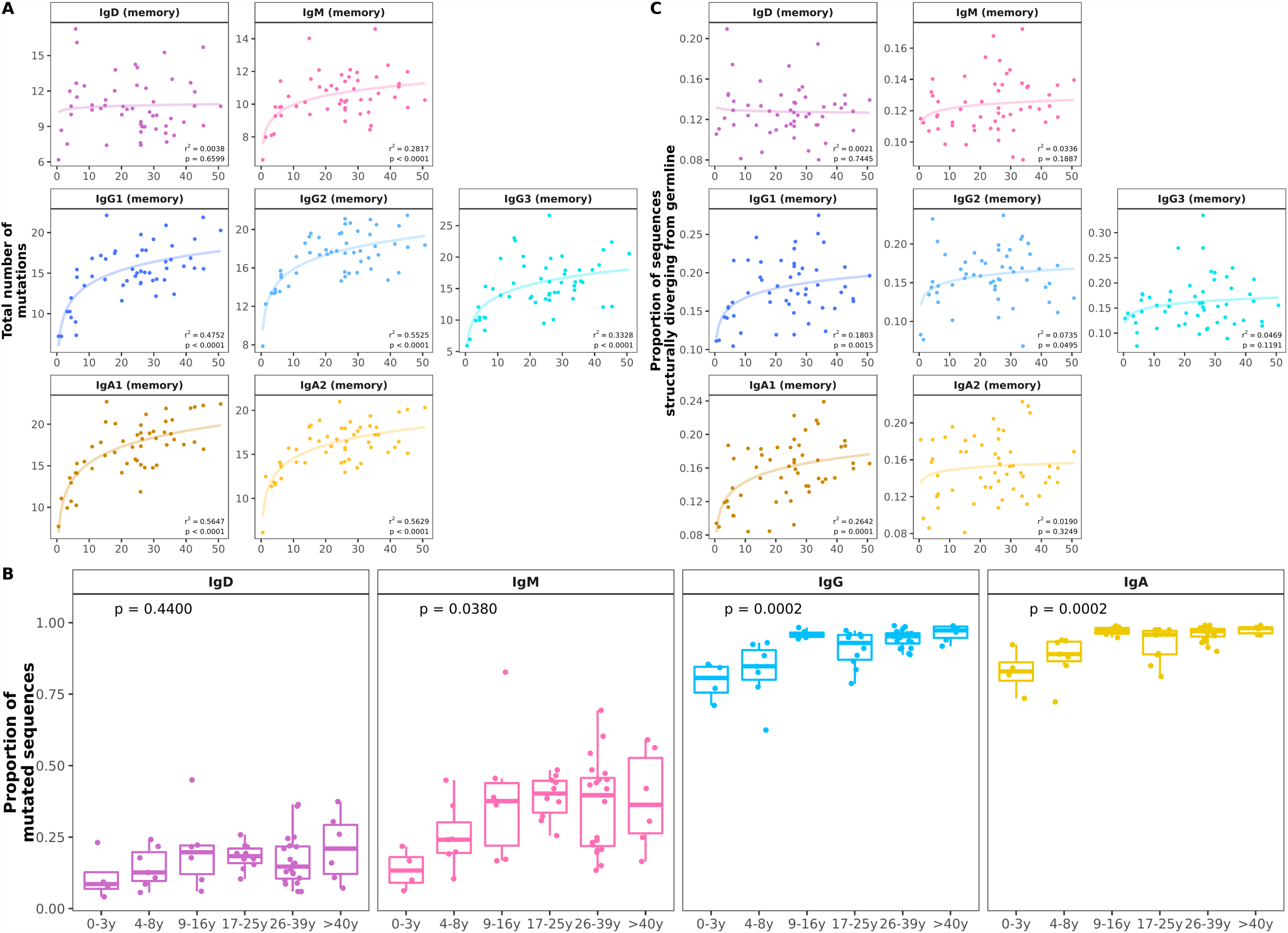
Age-related changes in somatic hypermutation and predicted antibody structure. *A* Mean number of V gene mutations by individual and B-cell subset with fitted logarithmic curves. Somatic hypermutation increased mainly in the first 10 years of life with some differences between cell subsets. *B* The proportion of memory IgD and memory IgM out of all IgD/IgM antibody and the proportion of mutated IgG and IgA antibody within repertoires showed significant increases in the first 10 years of life. *C* The proportion of sequences structurally different from germline increased in early childhood in all B-cell subsets. Statistical differences between groups were tested using the Kruskal-Wallis test.

### V family and J gene usages change with age

Although previous work has observed common patterns of gene segment usage and has suggested a strong dependence on an individual’s germline genetic background (33, 34), the relative contributions to variance from age remained unclear. Proportions of sequences assigned to the different V gene families and J genes were calculated for each sample and B-cell subset. The overall distribution of V family and J gene usage were different in older individuals compared with younger age groups. In particular, frequencies of V1 family sequences significantly decreased with age in naïve and mutated IgD and IgM sequences. This decrease was also observed in IgG and IgA antibody although with higher individual variation in older age groups (**Figure 1A**). No clear pattern was found in the usage of the other V families by age (**Supplementary figure 3A**). Such changes in V1 family genes were due to age-related alterations in several V genes, particularly VH1-8 (**Supplementary figure 5**).

**Figure 3:**
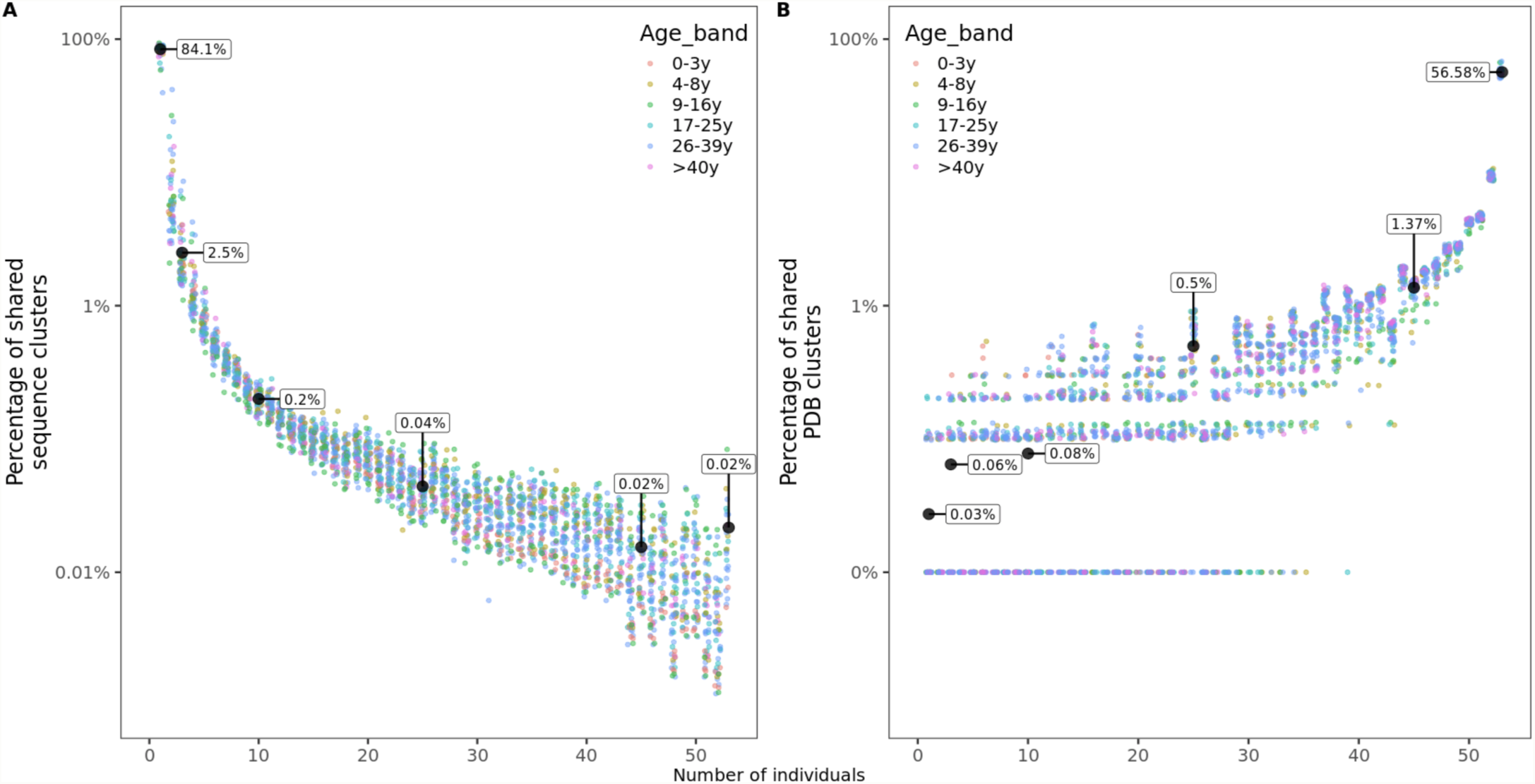
Sharing of sequence and structural clusters among the 53 healthy participants of different ages. *A* Percentage of sequence clusters shared by n individuals. *B* Percentage of structural clusters shared by n individuals. For structural clusters, zeros were replaced by 0.01% to be displayed on a logarithmic scale but labeled as 0%.

**Figure 4:**
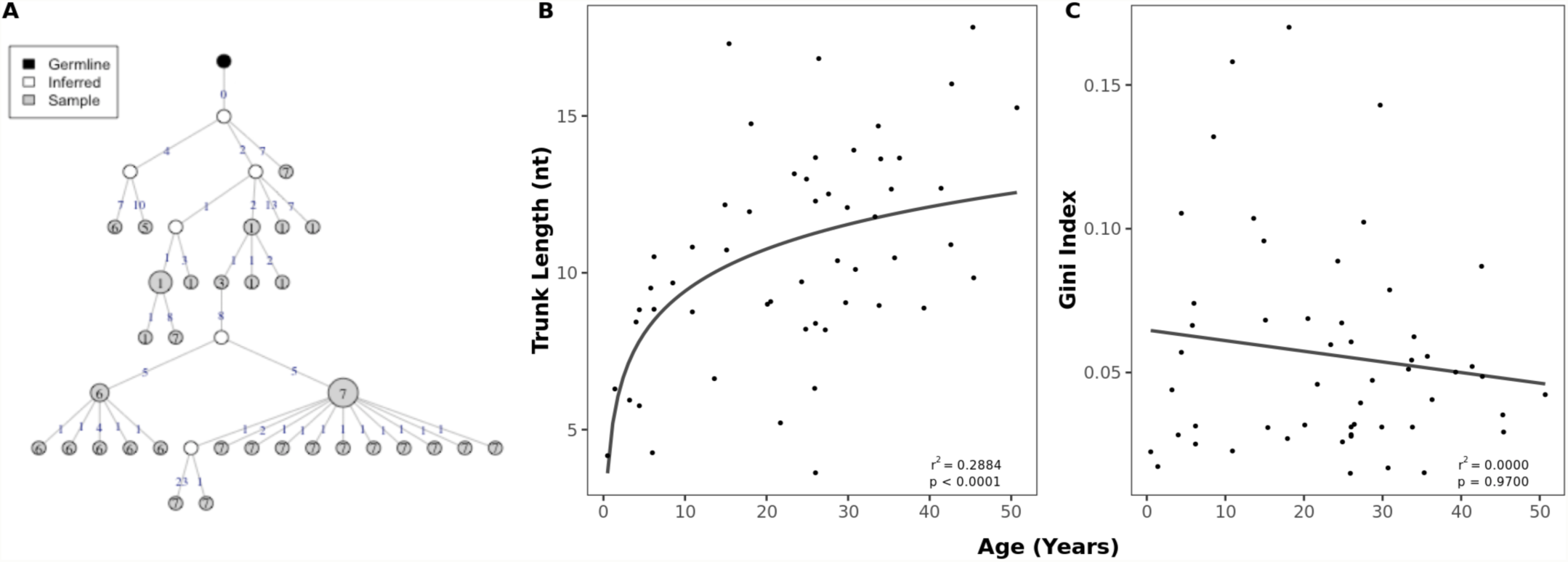
Age-related changes in clonal expansions. *A* Example lineage tree with each node representing a sequence and the size of the node indicating the number of identical sequences. The number of mutations between the sequences (nodes) is shown on top of the connecting lines. *B* Correlation between age and mean trunk length with a fitted logarithmic curve. *C* Correlation between mean Gini index and age with a fitted linear model.

**Figure 5:**
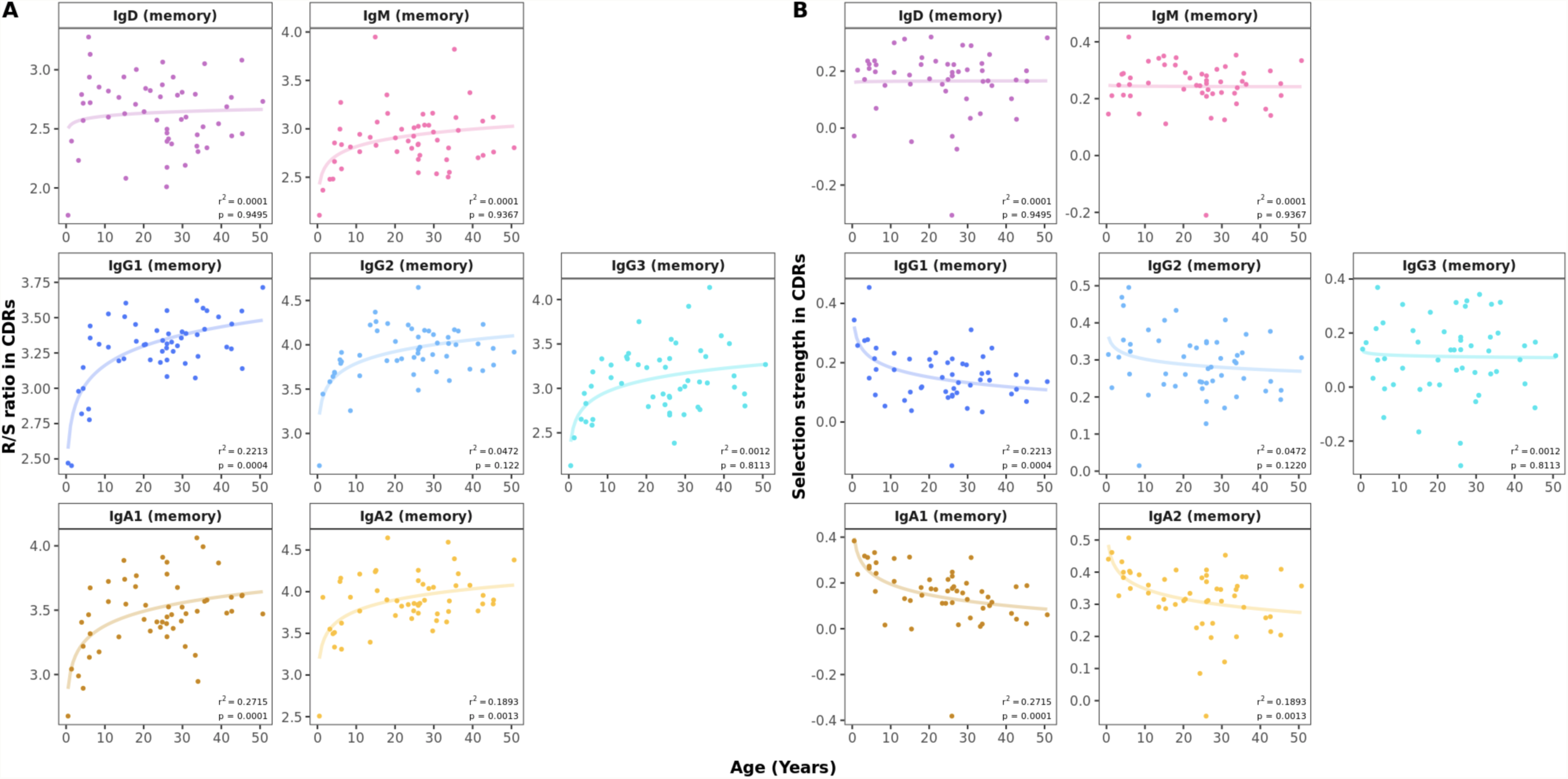
Age-related changes in antigen-driven selection. **A)** Mean R/S ratio in V gene CDRs as a measure of selection pressure showed an increase in early childhood in all B-cell subsets. For sequences with replacement but no silent mutations, the number of silent mutations was set to 1. **B)** Mean selection strength in CDRs calculated using BASELINe decreases with age in class switched subsets.

There were also changes in the overall J gene usage over the first 10 years of life marked by a significant decrease in the frequencies of sequences assigned to J6 in IgG antibody (**Figure 1B**). Frequencies of the other J genes by age group are shown in **Supplementary figure 3B**. In line with previous work (35, 36), we find that BCR sequences with rearranged J6 gene have longer junctions (**Supplementary figure 3C**). Along with a declining J6 usage with age, a significant decrease in junction length was observed in IgG subsets of older individuals (**Figure 1C**). However, even within IgG J6 antibody, junction length significantly decreased with age indicating that shorter junctions in older individuals are not solely the result of altered J gene usage (**Supplementary figure 4**).

### Somatic hypermutation exponentially increases in the first 10 years of life

There was a significant increase in SHM in all antigen-experienced subsets with age, which was most prominent in the first 10 years of life (**Figure 2A**). Substantial changes in mutation counts were found in all IgA and IgG subsets with exponential increases in children under 10 years and more linear progression between 10 and 50 years. IgM memory showed the smallest change of all subsets with some increase in children and a plateau from the 2^nd^ decade while there was no age-dependent change for IgD memory antibody. However, the proportion of mutated IgM antibody per sample increased from 0.1 in 0-3 year olds to an average of 0.4 in older individuals (**Figure 2B**). An age-related increase in the proportion of mutated sequences was also seen for IgA and IgG although at a higher level (**Figure 2B**).

### Sequences with predicted antibody structures diverging from germline increase with age

Crystallographic studies have shown that antibody CDR-H1 and CDR-H2 loops can adopt a very limited number of structural conformations, known as canonical loop classes (37, 38). These canonical classes are considered to be separate and distinct structures of the CDRs and can be rapidly and accurately annotated by SCALOP (23). The proportion of sequences in which either CDR-H1 and CDR-H2 had switched from the canonical class of their germline increased with age for most antigen-experienced subsets, similar to the increasing mutation number with age (**Figure 2C**).

Structures of CDR3 were predicted by mapping sequences to antibody structures in the PDB and annotated with a PDB code identifier. The proportion of every PDB cluster within individual and repertoire was calculated and normalized to zero mean and unit variance across individuals. PDB cluster usages were similar across individuals and age with a small number of positive outliers (frequent usage) that were private to each individual (**Supplementary figure 6**).

**Figure 6:**
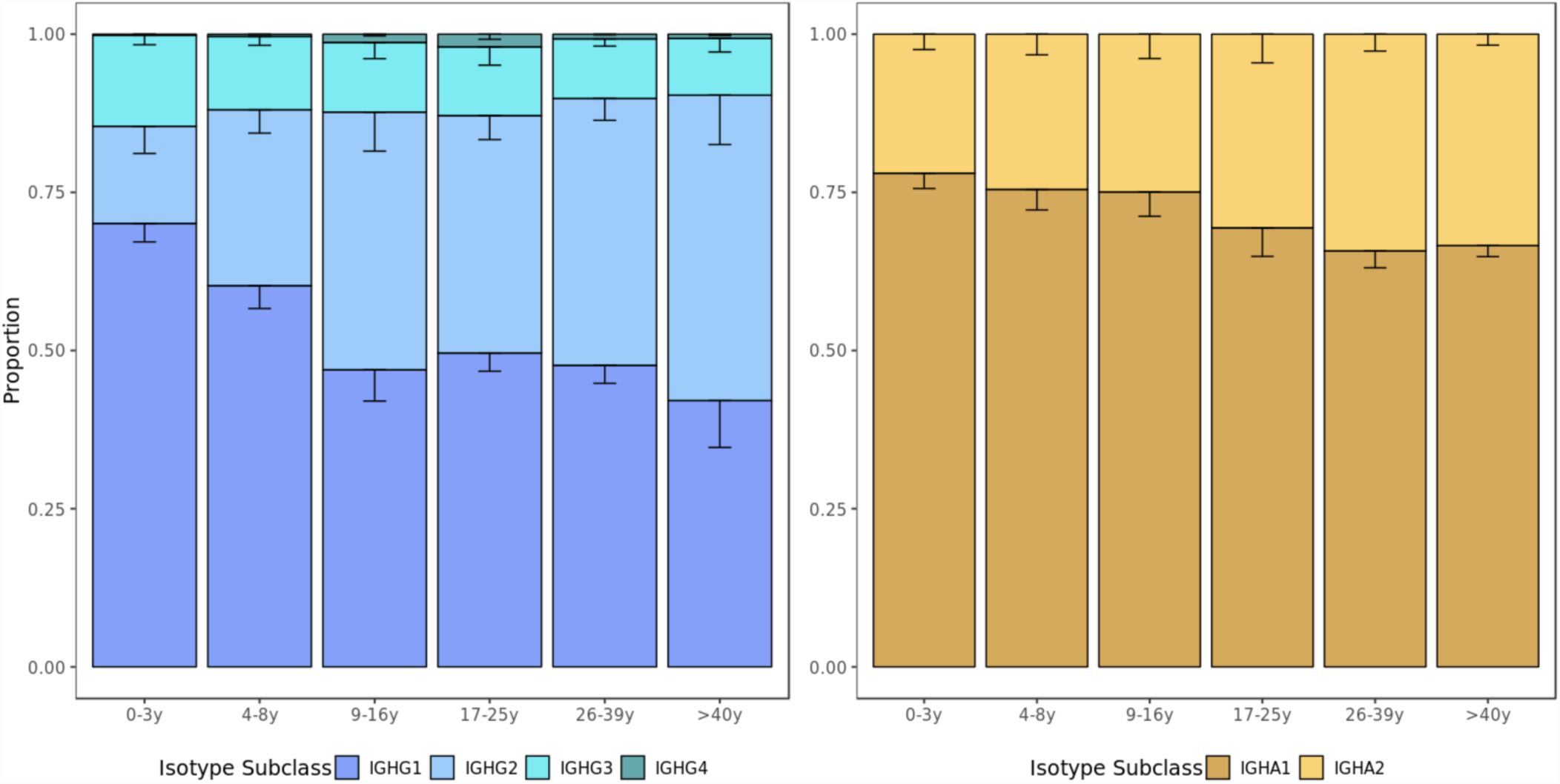
Usage of IgG and IgA subclasses by age group. The IgG and IgA isotype subclass usage changes with age. Error bars represent standard error of the mean.

### Structural but not sequence clusters are commonly shared between individuals

For each of the 53 individuals in this study, we calculated the frequency of sequence clusters (i.e. clonally related sequences) that are unique to the individual, the frequency of clusters that are shared with two, three or more subjects. Overlap with n subjects was quantified as the number of clusters shared with only n individuals divided by the total number of clusters in an individual’s repertoire. We found that on average, 84.1% of clusters were unique to the individual, while 2.5%, 0.2%, 0.04% and 0.02% of clusters were shared with 2, 10, 25 and 45 or more other individuals, respectively (**Figure 3A**). Sharing of structural clusters, however, was much more frequent with the majority of clusters (57%) shared by all 53 individuals and on average only 0.03% of clusters unique to the individual (**Figure 3B**). Neither sequence nor structural cluster sharing showed age-related changes.

### Older individuals display more mature clonal lineages and antibody with antigen-driven selection

Lineage trees were constructed from clusters of clonally related sequences and used to determine the evolutionary relationship within clusters (**Figure 4A**). The mean trunk length, representing the distance between the most recent common ancestor and germline sequence as a measure of the maturity of a lineage (39), greatly increased with age (**Figure 4B**). There was no relationship between age and the Gini index, which predicts whether lineages are dominated by a single clone (high index) or has a broad branching structure (low index) (**Figure 4C**). To account for differences in read depth, these characteristics were calculated on subsampled data so that the numbers of sequences were similar between individuals.

Insights into the process of antigen-driven selection can be gained by analyzing the mutational pattern in antigen-experienced repertoires. The R/S ratio in CDRs showed a marked increase in all antigen-experienced subsets between 0 and 10 years of life (**Figure 5A**). In samples from study participants older than 10 years, the R/S ratio was largely constant with values of around 3-3.5 in all B-cell subsets. In contrast, the R/S ratio was less variable and lower in FWRs compared with CDRs and no association with age was found (**Supplementary figure 7**). Next, we determined selection pressure using a Bayesian estimation of antigen-driven selection (BASELINe), which calculates selection by comparing the observed mutations to expected mutations derived from an underlying SHM targeting model (18). In CDRs, there was a general trend towards an age-associated decrease in selection strength for IgA and IgG antibody whereas this was constant across age for IgD or IgM sequences (**Figure 5B**). The statistical framework used to test for selection was CDR_R / (CDR_R + CDR_S), which normalizes for the observed increase in the total number of mutations with age.

**Figure 7:**
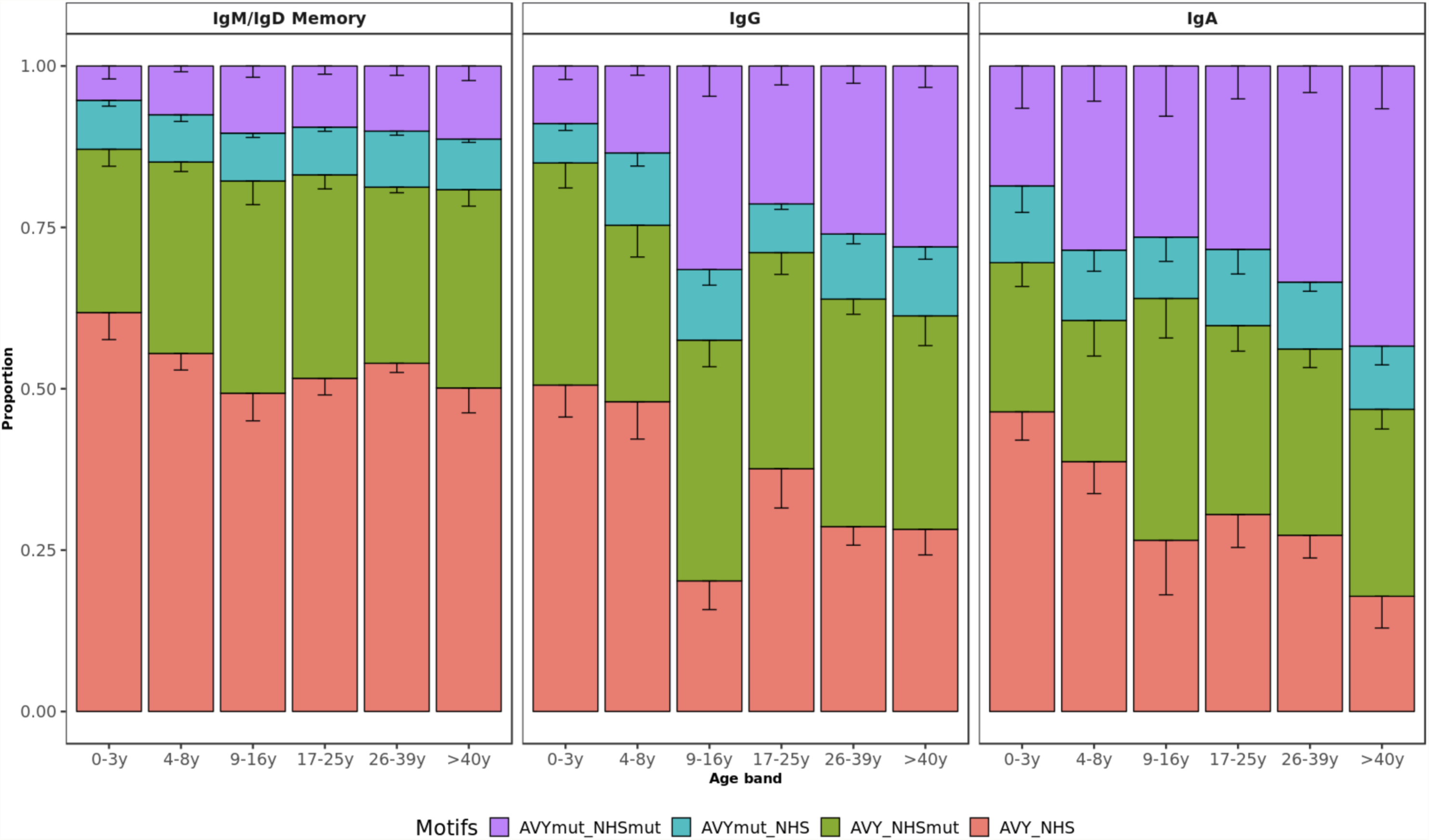
VH4-34 motifs by age group. Bar plots represent the proportion of sequences with mutated AVY and/or NHS motifs in IgD/IgM, IgG and IgA. Error bars indicate standard error of the mean. Proportion of sequences with both unmutated motifs decreases with age.

### Usage of IgG2 and IgA2 subclasses increase with age

Subclass usages were calculated within IgA and IgG repertoires to explore age-dependent class-switching patterns. In most age groups, IgG1 sequences were the most commonly detected, followed by IgG2, IgG3, and IgG4 sequences. However, the proportion of IgG2 sequences increased with age (p=0.0140, Kruskal-Wallis by age group) at the expense of lower usage of IgG1 (p=0.0086, Kruskal-Wallis) and IgG3 (p=0.1900, Kruskal-Wallis) sequences in older individuals. Similarly, IgA1 was most commonly used in all age groups and there was a non-significant trend towards a higher proportion of IgA2 sequences with age (p=0.0960, Kruskal-Wallis) (**Figure 6**).

### Repertoires from older individuals contain more self-tolerant sequences

Self-reactive antibodies share sequence characteristics that can be explored by Ig-seq. These include an increased usage of certain V genes, mainly VH4-34, and usage of longer CDR3 with positively charged or hydrophobic residues (40–42). We investigated how these metrics vary with age in healthy individuals. Apart from the decreasing junction length in IgG subsets (**Figure 1C**), we found that age has no impact on charge or hydrophobicity of BCR repertoires (**Supplementary figure 8**). VH4-34 usage was also unrelated to age whereas a more detailed SHM analysis including self-reactive motifs of VH4-34 sequences revealed an age-specific pattern. The VH4-34 germline contains an Ala-Val-Tyr (AVY) hydrophobic patch in FWR1 that is not present in other V segments and is thought to contribute to the self-reactive property of this gene (43, 44). Another feature of the VH4-34 germline associated with autoimmunity is the presence of an Asn-X-Ser N-glycosylation sequon (NHS) in CDR2 that modulates antibody avidity (45). Previous research has shown that mutating one or both of these motifs drives specificity of these sequences away from self, thereby contributing to peripheral tolerance. Lower frequencies of both unmutated AVY and NHS were present in healthy elderly individuals while there was a relative accumulation of single and double-mutated motifs in VH4-34 with age (**Figure 7**). This pattern was observed across all antigen-experienced subsets but was only statistically significant for IgA and IgG antibody (p=0.0110 and p=0.0036 respectively; p=0.1800 for IgM/IgD memory; Kruskal-Wallis test).

**Figure 8.**
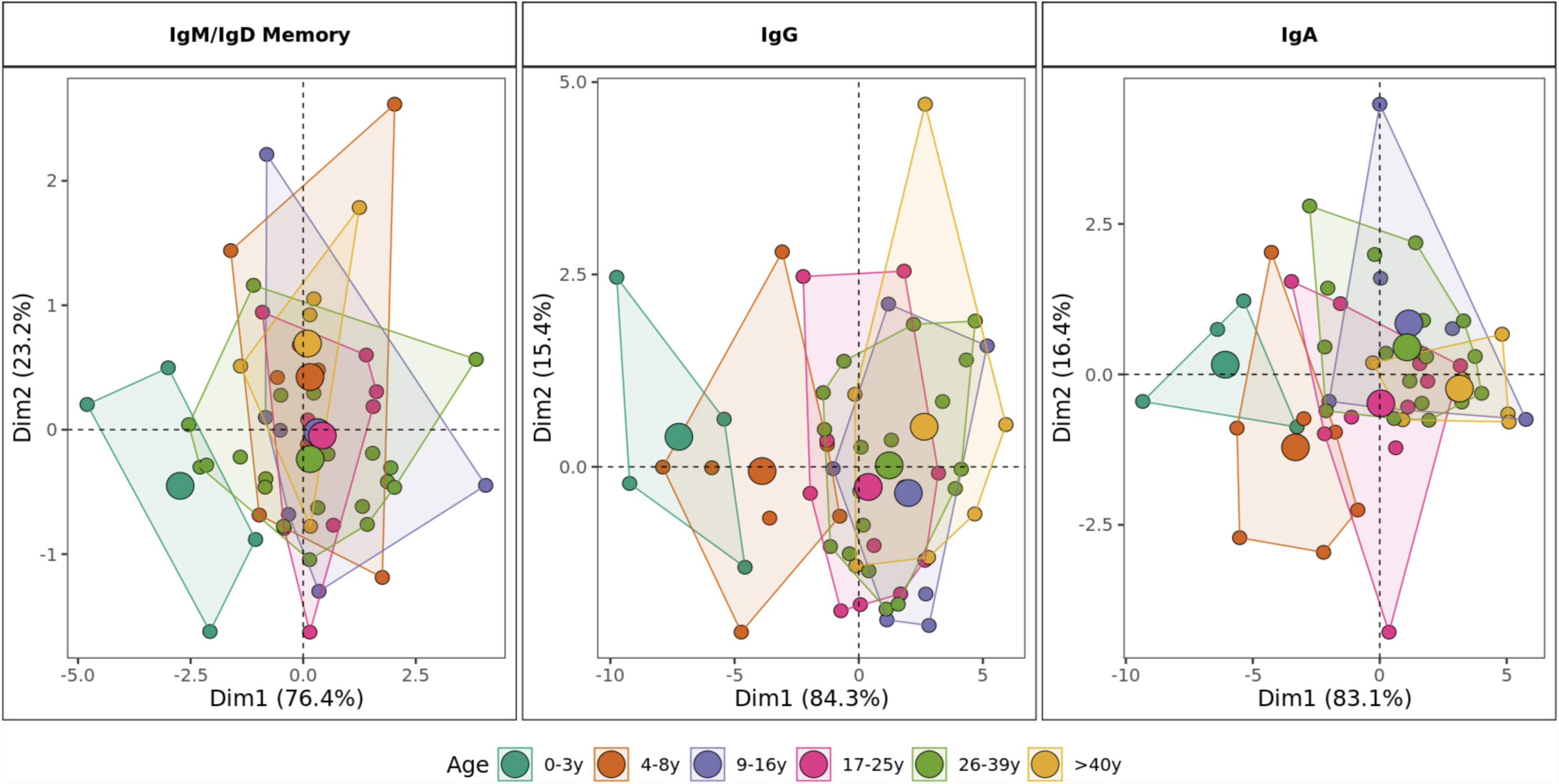
Stratification of BCR repertoires by age group. Principal component analysis by age category including mutation rate, R/S ratio, V1 gene family usage, J6 gene usage, junction length and proportion of sequences structurally divergent from germline as variables. For class-switched IgG and IgA, the proportion of IgG2 and IgA1 are included respectively. Areas are the convex hulls of the age group and the largest point of one color represents the center of that hull.

### Combining age-related repertoire features distinguishes between children and adults

Principal component analysis (PCA) based on the age-driven variables including mutation, R/S ratio, junction length, gene usage and proportion of sequences structurally divergent from germline clearly showed distinct grouping of children younger than 9 years old and individuals older than 10 years old in antigen experienced repertoires. This distinction was most clearly observed in the class-switched IgG and IgA repertoires. In IgD/IgM mutated sequences, children less than three years old were separate from other individuals whereas the repertoire characteristics in older age categories overlapped.

## Discussion

In this study, we found an extensive maturation of B-cell responses in the first 10 years of life consistent with what would be expected with cumulative antigen exposure and a generally more developed and stable B-cell compartment in older individuals. Further antibody repertoire alterations continue to be made thereafter, although at a lower rate. The results presented here constitute the most in-depth evaluation of the BCR repertoire with age. This study also provides a detailed reference data set of isotype and subclass-specific BCR repertoires of healthy individuals across a relevant age range and stresses the importance of using well-selected, age-appropriate controls in future studies.

Previous studies have suggested that immunoglobulin gene usage is strongly genetically determined as it was conserved between monozygotic twins and across multiple time points within a given individual (7, 33). We found age-dependent alterations in both V family and J gene usage in antigen-experienced repertoires suggesting either polyclonal negative selection of V1- and J6-containing B cells or positive selection of non-V1/J6-bearing B cells during maturation of the adaptive immune system. However, here we also saw that V family gene usage changed in naïve repertoires that are supposedly unaffected by antigen exposure and not subject to antigen-driven selection pressure, indicating preferential development and/or survival of V1-bearing B cells in young children. Although the potential benefit and mechanism behind these age-related V family gene alterations remain unclear, these findings suggest that immunoglobulin gene usage in developing B cells is less conserved than previously thought.

In line with earlier findings (11, 46, 47), we observed extensive maturation of antigen-experienced repertoires characterized by accumulation of somatically hypermutated B-cell antibody with evidence of strong positive selection in older individuals. The observed decrease in selection pressure in some class-switched subsets indicates that young individuals show accelerated dynamics to achieve highly selected sequences compared with older individuals. Of note, detailed analysis allowed to investigate characteristics of mutated IgM/D antibody separately, which were observed at a higher frequency and with a greater number of mutations in older individuals. These findings indicate that the pool of circulating peripheral blood naïve B cells is continuously diminishing with age, possibly contributing to a decreasing capacity to effectively respond to novel antigens in older individuals (48). We also observed a substantially higher proportion of unmutated IgA/G antibody in young children compared with adults (49), a finding that has not yet been recognized and is of unknown significance. However, these results are in line with previous *in vitro* studies (50) demonstrating that class-switch recombination and somatic hypermutation can occur independently and suggest class-switching to be an important element of B-cell responses in young children.

Along with other characteristics indicative of antigen-driven maturation we found that the proportion of sequences with structures differing from germline increased with age, which was most pronounced for IgG1 and IgA1 subsets. To date, there is limited information on predicted antibody structures derived from high-throughput adaptive immune receptor repertoire sequencing data (51, 52). In line with measures of antigen-driven selection, there was a positive linear relationship between number of mutations and structural alterations of antigen-experienced sequences indicating that alteration of the three-dimensional structure is important to achieve high specificity and affinity of the antibody. By annotating individual sequences with PDB codes, we were able to investigate commonalities of CDR3 structures between individuals. In particular, in contrast to sharing on the sequence level, the majority of PDB clusters were public while only a very small percentage of PDB clusters were private to the individual. Although this comparison is influenced by the much smaller number of potential PDB clusters, the use of common PDB clusters indicates that a large number of different sequences can underlie similar antibody structures. Future work, such as the investigation of PDB usage in patients with immune disorders, will help determine how antibody structures can be used to assess global immune responses.

We found an increase in the usage of IgA2/IgG2 antibody with age, similar to what has been seen in a recent study on the isotype subclasses surface expression of peripheral blood B cells (53). While human IgG subclasses have been extensively studied (54), there is limited information on the functional difference between the two IgA subclasses, whose structures mainly differ in the length of the hinge region (55). IgG2 has been implicated in the immune responses to capsular polysaccharides of bacteria such as *S. pneumoniae* that are commonly colonizing the oropharynx of young children and thereby induce polysaccharide-specific serum antibody (56). Our findings also match the sequential model proposed for CSR: with age, and after multiple encounter with the same antigen, class-switched memory B cells re-enter the germinal center to undergo a second round of CSR and switch towards more downstream constant region genes (57).

The majority of early immature human B cells display self-reactivity and although most of these are removed during B-cell development, a substantial proportion of mature B cells may still be directed against autoantigens (40). Antibodies encoded by germline VH4-34 are intrinsically self-reactive antibodies mediated by a hydrophobic patch and a glycosylation sequon (43, 45). Unmutated VH4-34 antibody are more common in naïve than antigen-experienced repertoires as receptor editing of these antibodies drives specificity away from self (44, 58). In contrast to adults, we found that a substantial proportion of VH4-34 IgG and IgA antibody from children are unmutated, with frequencies gradually decreasing with age. Previous work has shown that germline VH4-34-expressing IgG B cells recognized antigens from commensal gut bacterial (58) and hence, the higher frequency of these cells in children may be related to ongoing immune responses against gut pathogens in this age group.

This study used Ig-seq technology coupled with bioinformatic methods to study in detail the BCR repertoires of healthy individuals and investigate the effect of age on repertoire characteristics. We chose a cross-sectional study design and – although unlikely – can therefore not exclude that longitudinal assessment of maturation on an individual basis may differ from the presented findings. For practical reasons, the number of input cells was variable between study participants, which resulted in variable sequence numbers per sample. For analysis where sequence number variability was considered to be of major relevance, such as constructing lineage trees, subsampling to an equal number of sequences per individual was performed.

We were able to map in detail the characteristics, magnitude and speed of age-dependent maturation of BCR repertoires. Combining age-related variables using a PCA allowed clear separation of individuals younger than 10 years from older study participants, which was most pronounced in IgG repertoires. Our analysis now allows comparisons to be made in the BCR repertoires of healthy individuals to patients with altered immune states such as primary or secondary immunodeficiency (4) or infectious disease (59, 60). By elucidating patterns that are associated with cumulative antigen exposure and an evolving immune system, this research offers important insight into adaptive immune system responses in humans. The mechanisms behind the development of clinically relevant autoimmunity is still poorly understood and the findings in this study show a substantial intrinsic capacity to produce self-reactive B cells, which may be essential to achieve the diversity needed for the defense against commensal pathogens in early life.

In summary, by studying the maturation of the healthy BCR repertoire with age, we found characteristics indicative of a maturing B-cell system consisting of alterations in immunoglobulin gene usage, increased levels of SHM associated with strong positive selection, and canonical class usage that differed considerably from germline structures. Repertoires from older individuals more frequently contained antibody using more downstream constant region genes that are involved in the immune response to polysaccharide antigens. With accumulating mutations, germline-encoded self-reactive antibody were seen less with advancing age indicating a possible beneficial role of self-reactive B-cells in the developing immune system. This study provides a reference data set of isotype subclass-specific BCR repertoires and stresses the importance of using well-selected, age-appropriate controls in future studies.

## Supplementary information

**Supplementary figure 1.**
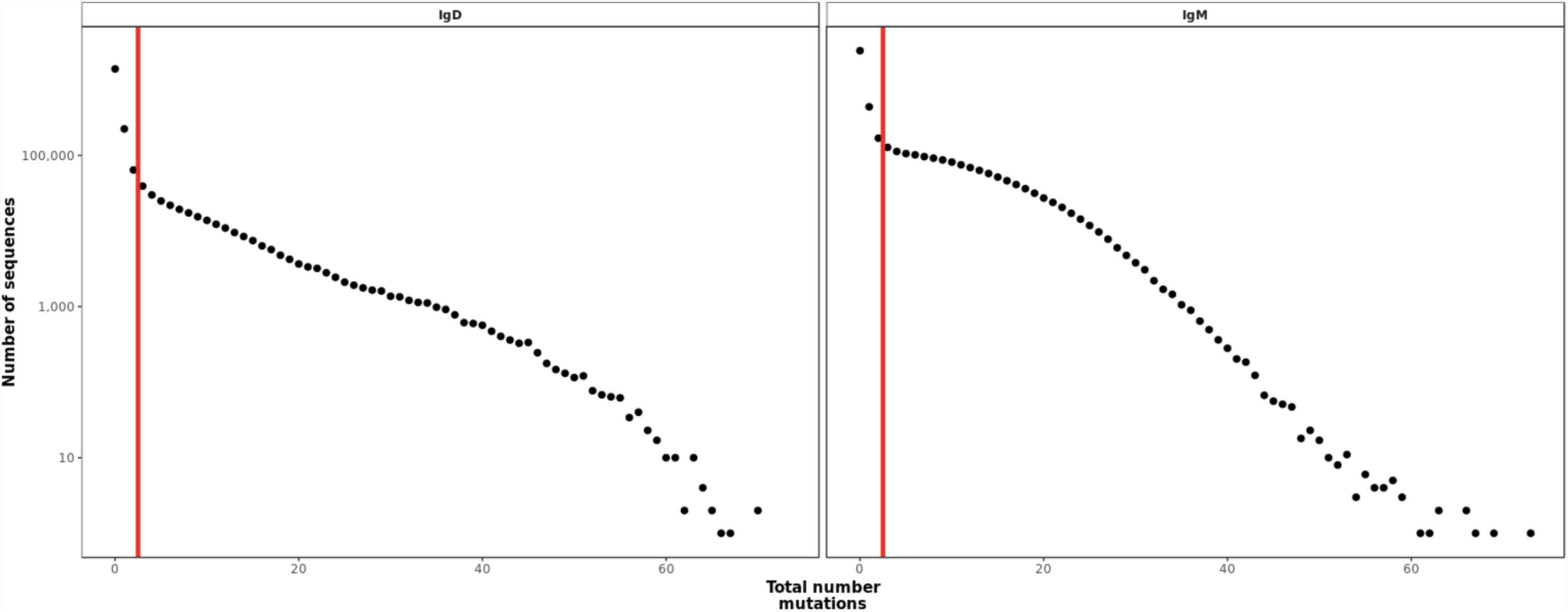
Distribution of somatic hypermutation in all IgD and IgM transcripts. The vertical line indicates the threshold chosen (mutation n between 2 and 3) to separate naïve and memory repertoires for IgM and IgD sequences.

**Supplementary figure 2:**
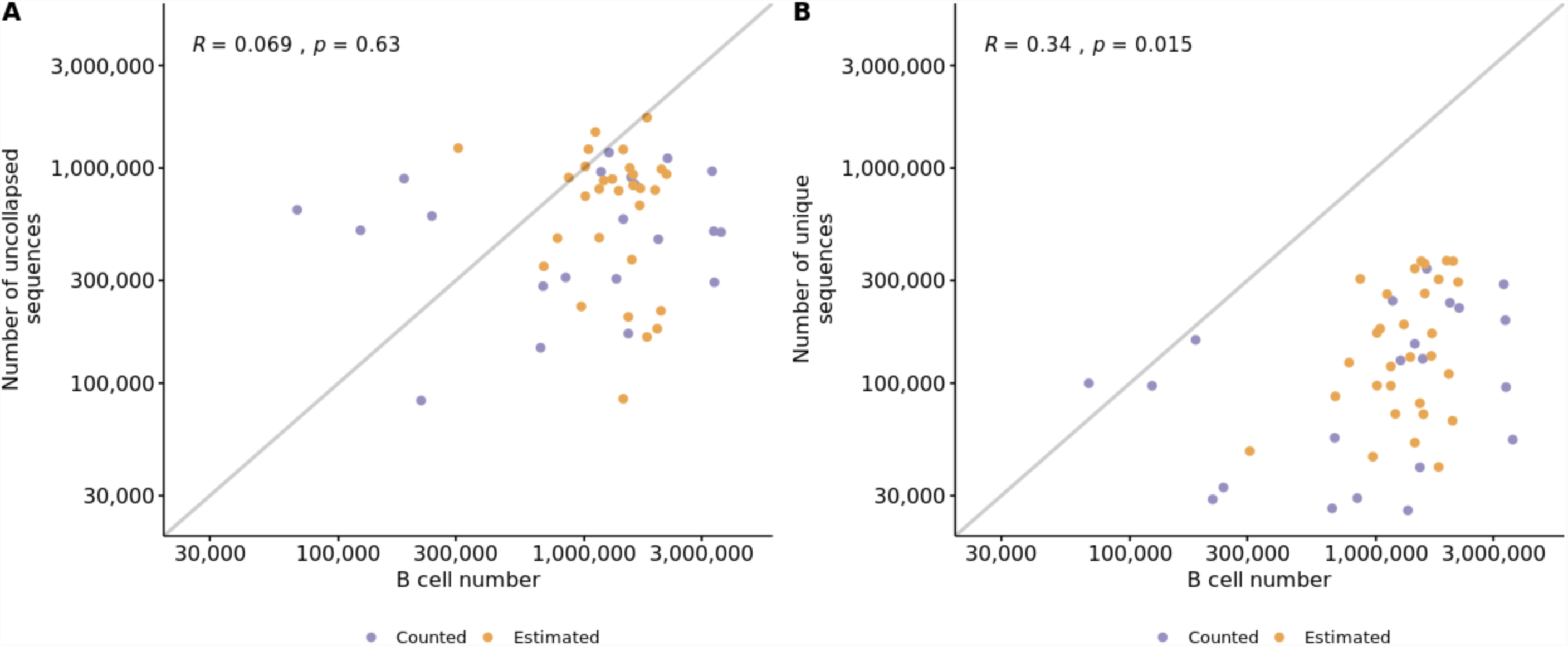
Correlation between cell number in a sample, and the number of sequences for that sample *A* before and *B* after collapsing (Pearson correlation coefficient). The B-cell number was either based on actual counts or estimated using PBMC counts and the median percentage of age-dependent reference values.

**Supplementary figure 3.**
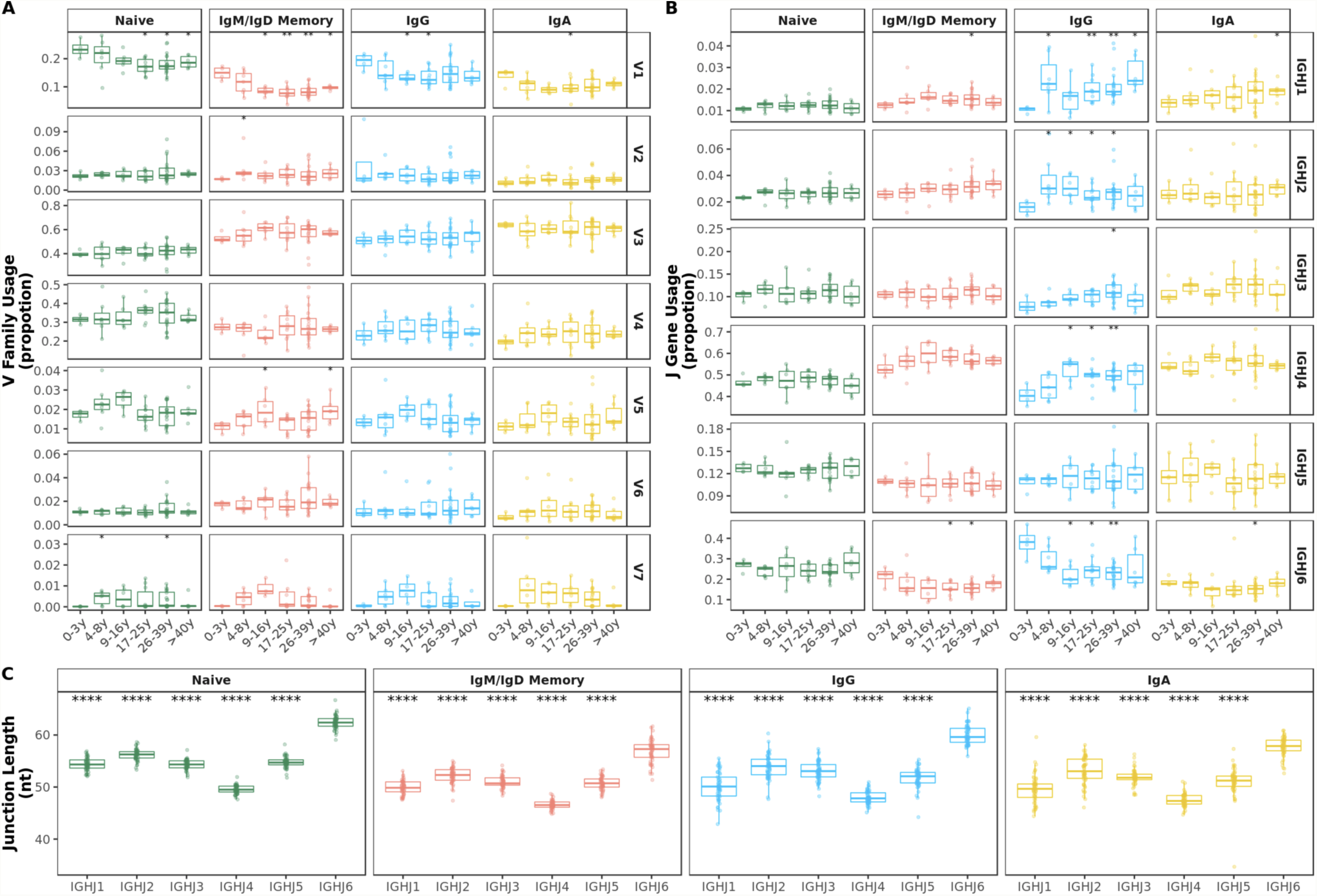
*A* V family and *B* J gene usage by age band. Comparison of each age group to the 0-3y group was performed using the Wilcoxon test. *C* IGHJ6 transcripts show significantly longer junctions. Comparison of each gene to IGHJ6 was performed using the Wilcoxon test. *p<0.05, **p<0.01, ***p<0.001, ****p<0.0001

**Supplementary Figure 4:**
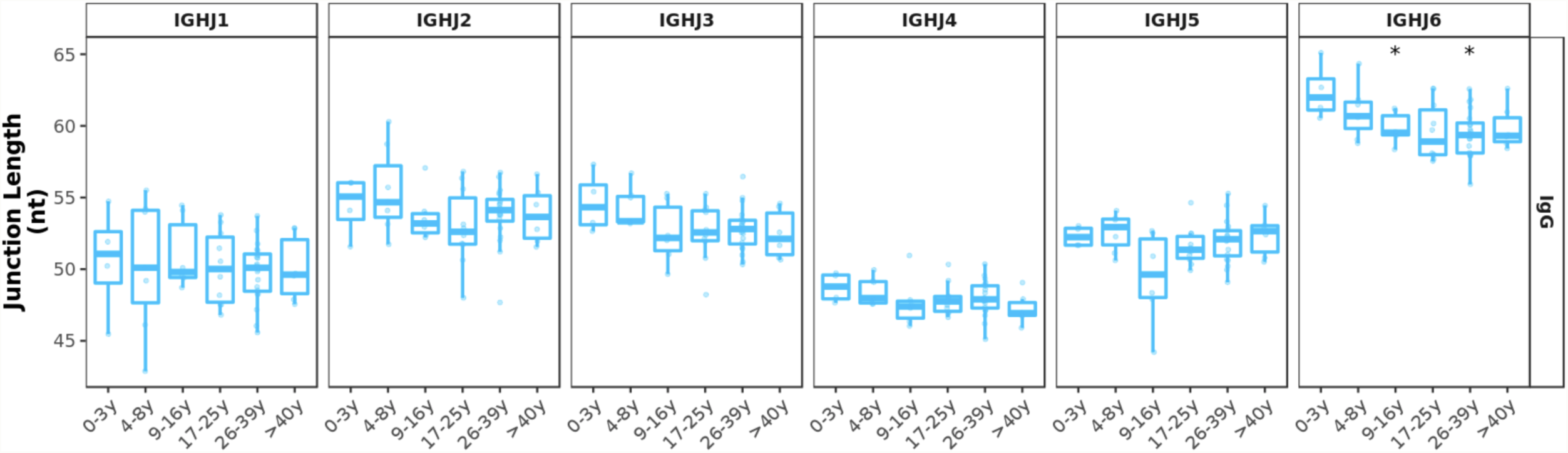
Junction length decrease in IgG transcripts is still apparent within J gene and is only significant in transcripts with IGHJ6. Comparison of each age group to the 0-3y group was performed using the Wilcoxon test. *p<0.05

**Supplementary figure 5.**
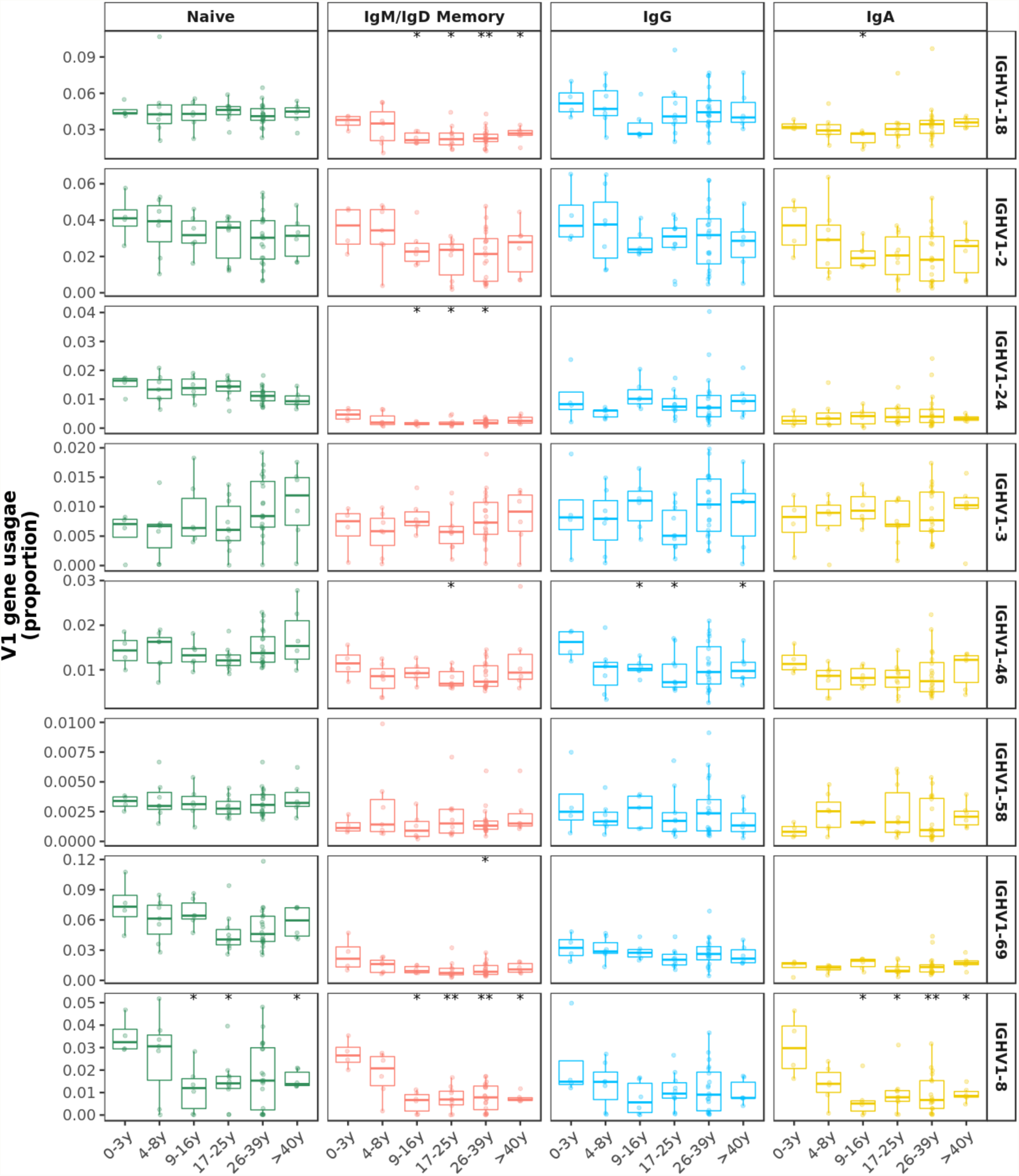
Proportion of the top 8 V1 family genes by age band. The decrease seen in V1 family usage is a result of a decrease in multiple individual genes. Comparison of each age group to the 0-3y group was performed using the Wilcoxon test. *p<0.05, **p<0.01

**Supplementary Figure 6.**
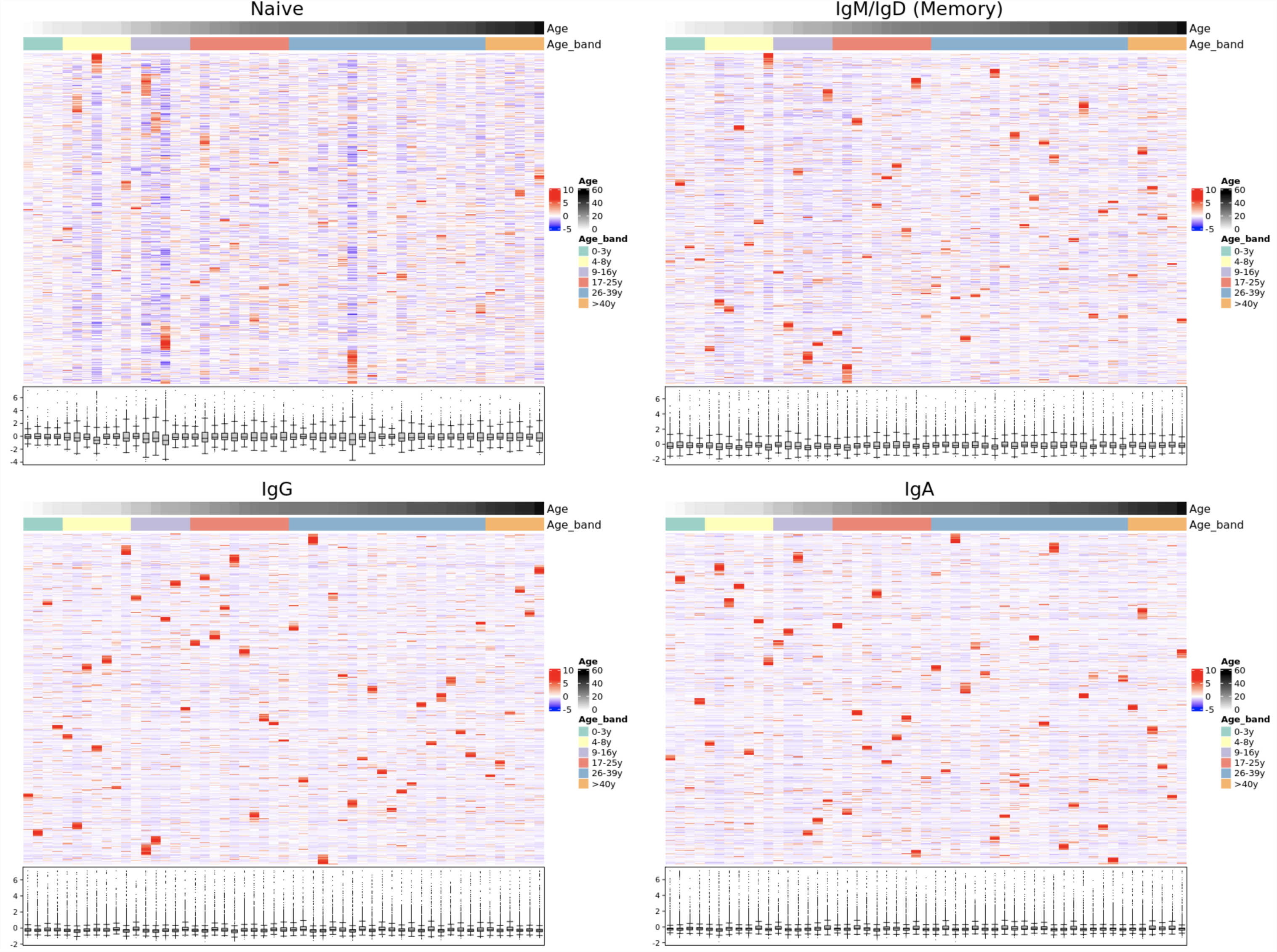
The structural composition of the naïve, IgM/IgD memory and class-switched IgG and IgA repertoires. Heatmaps are normalized by row which represents usage of one PDB cluster. Distributions of the normalized PDB cluster usage by individual is shown as a boxplot. Samples (columns) are ordered by age and PDB clusters (rows) are hierarchically clustered.

**Supplementary figure 7.**
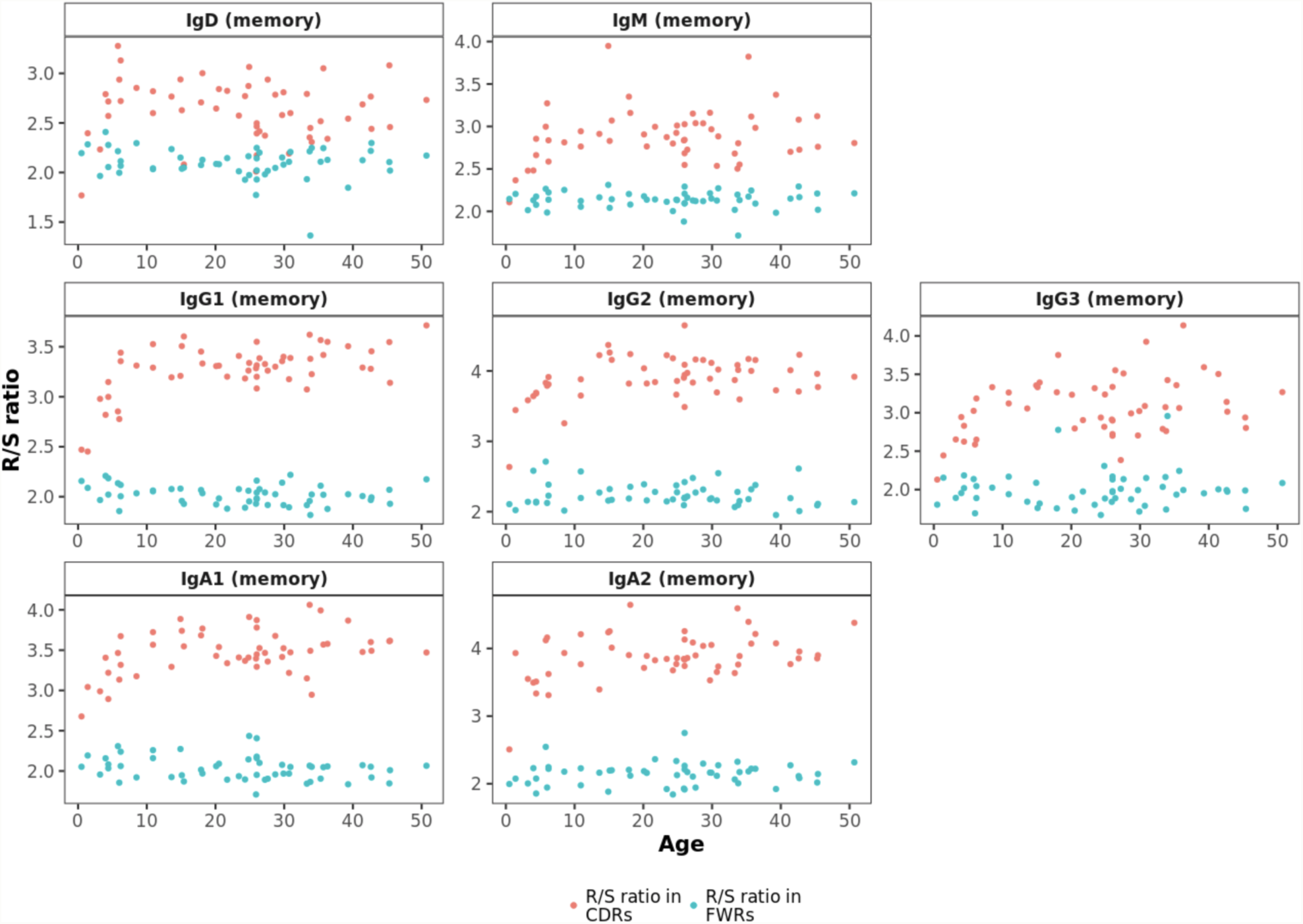
R/S ratio in FWRs does not correlate with age and is lower compared with CDRs.

**Supplementary figure 8.**
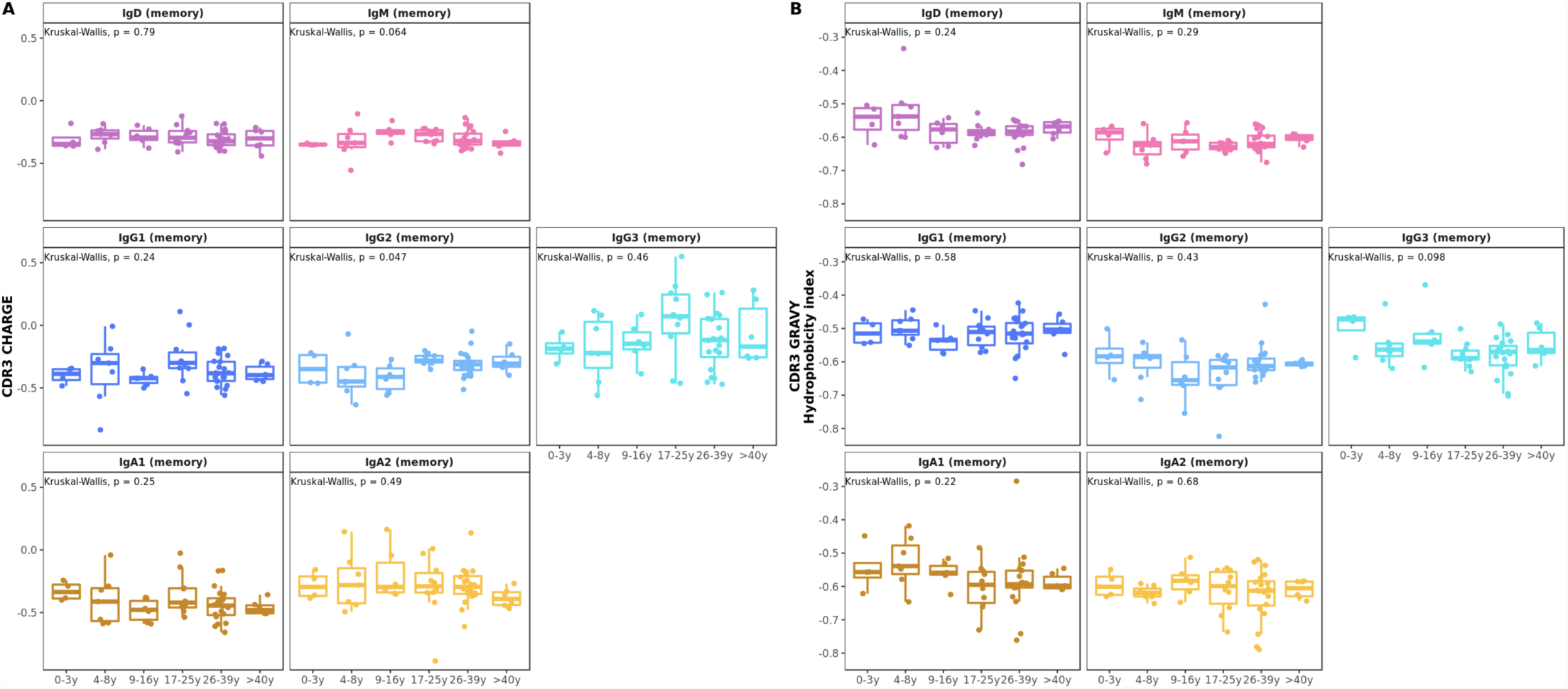
*A* CDR3 charge and *B* CDR3 hydrophobicity index do not correlate with age in healthy controls.

**Supplementary table 1.**

| Participant | Sex | Age (y) | Cell type | PBMC number | B cell number | B cells estimated (E) or counted (C) | RT PCR | Raw Sequences | Unique Productive Sequences | IgD (naïve) | IgM (naïve) | IgD (memory) | IgM (memory) | IgG1 (memory) | IgG2 (memory) | IgG3 (memory) | IgG4 (Memory) | IgA1 (Memory) | IgA2 (memory) | IgE (memory) |
| --- | --- | --- | --- | --- | --- | --- | --- | --- | --- | --- | --- | --- | --- | --- | --- | --- | --- | --- | --- | --- |
| HC_1 | F | 0.5 | PBMCs | 14400000 | 3024000 | E | AEG,MD | 1904663 | 285839 | 72572 | 159399 | 2240 | 7963 | 21480 | 4276 | 4875 | 38 | 9509 | 3429 | 58 |
| HC_2 | F | 1.4 | PBMCs | 6900000 | 1932000 | E | AEG,MD | 2326250 | 239052 | 62659 | 132670 | 5230 | 11630 | 16237 | 1263 | 3647 | 18 | 4585 | 1111 | 2 |
| HC_3 | M | 3.2 | PBMCs | 9900000 | 2376000 | E | AEG,MD | 1622639 | 221134 | 52374 | 115799 | 3386 | 18607 | 11164 | 4741 | 1819 | 54 | 9894 | 3246 | 50 |
| HC_4 | F | 3.9 | PBMCs | 9450000 | 2268000 | E | AEG,MD | 1571312 | 337616 | 55542 | 156394 | 14433 | 35134 | 35350 | 6421 | 7459 | 46 | 22599 | 4230 | 8 |
| HC_5 | F | 4.4 | PBMCs | 4200000 | 1008000 | E | AEG,MD | 1975420 | 93299 | 20948 | 44505 | 2070 | 10982 | 5957 | 3788 | 1587 | 42 | 2099 | 1280 | 41 |
| HC_6 | M | 4.4 | PBMCs | 1140000 | 273600 | E | AEG,MD | 1216544 | 28674 | 13955 | 9161 | 1689 | 1786 | 872 | 341 | 122 | 4 | 599 | 141 | 4 |
| HC_7 | F | 5.8 | PBMCs | 11100000 | 1998000 | E | AEG,MD | 1973589 | 128326 | 35276 | 44614 | 10339 | 9463 | 13050 | 5047 | 1526 | 53 | 7791 | 1164 | 3 |
| HC_8 | M | 6.0 | PBMCs | 12600000 | 2268000 | E | AEG,MD | 1818496 | 124567 | 7399 | 32174 | 1387 | 22205 | 37668 | 5395 | 5989 | 34 | 10097 | 2147 | 72 |
| HC_9 | M | 6.2 | PBMCs | 6150000 | 1107000 | E | AEG,MD | 2474481 | 255693 | 84048 | 129241 | 4096 | 12182 | 5861 | 4610 | 1268 | 46 | 9844 | 4117 | 380 |
| HC_10 | F | 6.2 | PBMCs | 10800000 | 1944000 | E | AEG,MD | 1337214 | 368861 | 29803 | 112882 | 7228 | 54220 | 42616 | 28527 | 7659 | 534 | 59361 | 25936 | 95 |
| HC_11 | M | 8.5 | PBMCs | 10800000 | 1944000 | E | AEG,MD | 880759 | 40494 | 18417 | 10920 | 1431 | 3099 | 2283 | 841 | 725 | 19 | 2126 | 633 | 0 |
| HC_12 | F | 10.9 | PBMCs | 12900000 | 2064000 | E | AEG,MD | 1802230 | 366617 | 65222 | 148236 | 11720 | 72104 | 20972 | 10050 | 9851 | 427 | 19081 | 8746 | 208 |
| HC_13 | F | 10.9 | PBMCs | 7920000 | 1267200 | E | AEG,MD | 1168944 | 25242 | 6459 | 7505 | 1519 | 5248 | 1829 | 977 | 261 | 25 | 1057 | 275 | 87 |
| HC_14 | F | 13.6 | PBMCs | 4740000 | 758400 | E | AEG,MD | 872906 | 26006 | 8819 | 8659 | 2185 | 4714 | 278 | 644 | 70 | 5 | 450 | 173 | 9 |
| HC_15 | F | 14.9 | PBMCs | 1920000 | 307200 | E | AEG,MD | 2717449 | 44254 | 1632 | 3581 | 1259 | 17735 | 5991 | 3376 | 1053 | 238 | 5918 | 3464 | 7 |
| HC_16 | M | 15.1 | PBMCs | 14700000 | 2352000 | E | AEG,MD | 659493 | 95429 | 40827 | 36587 | 2228 | 6421 | 1635 | 2474 | 384 | 67 | 3639 | 1056 | 111 |
| HC_17 | F | 15.4 | PBMCs | NA | NA | NA | AEG,MD | 995764 | 289287 | 41886 | 118609 | 4719 | 24748 | 37623 | 27409 | 6542 | 1142 | 24002 | 2592 | 15 |
| HC_18 | F | 17.9 | PBMCs | 7200000 | 864000 | E | AEG,MD | 1682823 | 302272 | 26679 | 68526 | 4680 | 55458 | 37658 | 10783 | 12027 | 1252 | 77485 | 7704 | 20 |
| HC_19 | M | 18.1 | PBMCs | 8400000 | 1008000 | E | AEG,MD | 1011146 | 28798 | 12418 | 7307 | 1269 | 2209 | 1350 | 870 | 158 | 151 | 2248 | 818 | 0 |
| HC_20 | M | 20.1 | PBMCs | 13200000 | 1584000 | E | AEG,MD | 1433354 | 356105 | 42060 | 104452 | 5655 | 64801 | 39036 | 22756 | 4546 | 67 | 61075 | 11647 | 10 |
| HC_21 | M | 20.5 | PBMCs | 9600000 | 1152000 | E | AEG,MD | 945221 | 96859 | 25143 | 28963 | 6092 | 21561 | 5861 | 3223 | 381 | 684 | 3195 | 1695 | 61 |
| HC_22 | M | 21.7 | PBMCs | 15000000 | 1800000 | E | AEG,MD | 1140675 | 40591 | 21975 | 8263 | 4543 | 4454 | 294 | 178 | 244 | 1 | 477 | 160 | 2 |
| HC_23 | F | 23.4 | PBMCs | 9000000 | 1080000 | E | AEG,MD | 1423314 | 151168 | 68644 | 25308 | 10563 | 9876 | 7757 | 11844 | 1094 | 88 | 12146 | 3815 | 33 |
| HC_24 | F | 24.3 | CD19+ |  | 3600000 | C | AEG,MD | 1650432 | 53250 | 2584 | 27411 | 648 | 13631 | 2320 | 2099 | 628 | 26 | 2098 | 1803 | 2 |
| HC_25 | F | 24.8 | PBMCs | 17100000 | 2052000 | E | AEG,MD | 764310 | 66786 | 12784 | 24047 | 3933 | 17222 | 2397 | 1562 | 308 | 0 | 3145 | 1386 | 2 |
| HC_26 | F | 24.9 | PBMCs | 18000000 | 2160000 | E | AEG,MD | 1683395 | 288827 | 35736 | 91187 | 6432 | 63062 | 18078 | 28519 | 2636 | 2025 | 26499 | 14562 | 91 |
| HC_27 | M | 25.9 | PBMCs | 12000000 | 1440000 | E | Mix I | 593223 | 52933 | 7679 | 27223 | 1394 | 10655 | 2280 | 2011 | 432 | 17 | 509 | 728 | 5 |
| HC_28 | F | 26.0 | PBMCs | 14000000 | 1680000 | E | Mix I | 1146458 | 133861 | 20396 | 54984 | 3362 | 27783 | 10845 | 10135 | 1339 | 123 | 2477 | 2399 | 18 |
| HC_29 | F | 26.0 | CD19+ |  | 2000000 | C | AEG,MD | 1710083 | 235120 | 30835 | 145027 | 3509 | 38873 | 5427 | 3685 | 1335 | 17 | 4627 | 1783 | 2 |
| HC_30 | F | 26.0 | PBMCs | 3300000 | 185000 | C | AEG,MD | 2347978 | 158947 | 49263 | 63128 | 7051 | 20867 | 6055 | 4737 | 1279 | 99 | 5434 | 1033 | 1 |
| HC_31 | F | 26.0 | PBMCs | 1725000 | 68000 | C | AEG,MD | 1524836 | 99870 | 25370 | 29303 | 6741 | 23073 | 4476 | 3962 | 708 | 102 | 4847 | 1288 | 0 |
| HC_32 | F | 26.4 | PBMCs | 14100000 | 1692000 | E | AEG,MD | 1581380 | 168295 | 56857 | 51011 | 4527 | 10671 | 12723 | 6925 | 6070 | 230 | 13898 | 5288 | 95 |
| HC_33 | F | 27.2 | PBMCs | 13000000 | 1560000 | E | Mix I | 1217105 | 71742 | 8699 | 28815 | 1976 | 22157 | 4257 | 2197 | 1173 | 36 | 1479 | 953 | 0 |
| HC_34 | F | 27.6 | CD19+ | 11250000 | 240000 | C | AEG,MD | 941177 | 31596 | 2636 | 6664 | 1330 | 13308 | 1496 | 2727 | 208 | 52 | 2155 | 1020 | 0 |
| HC_35 | F | 28.7 | PBMCs | 8700000 | 1044000 | E | AEG,MD | 1824932 | 176403 | 17952 | 66414 | 4320 | 48242 | 11813 | 8105 | 1447 | 249 | 11565 | 6271 | 25 |
| HC_36 | F | 29.7 | PBMCs | 8100000 | 972000 | E | AEG,MD | 1103351 | 45312 | 8848 | 15726 | 817 | 10931 | 2324 | 1406 | 233 | 10 | 3726 | 1291 | 0 |
| HC_37 | M | 29.9 | PBMCs | 12000000 | 1440000 | E | AEG,MD | 1909727 | 338646 | 44381 | 139814 | 7439 | 91713 | 11746 | 23501 | 1585 | 425 | 8537 | 9479 | 26 |
| HC_38 | F | 30.7 | PBMCs | 12750000 | 1530000 | E | AEG,MD | 1877484 | 366879 | 106835 | 148305 | 5245 | 22404 | 13883 | 27944 | 3806 | 323 | 24438 | 13686 | 10 |
| HC_39 | M | 30.9 | PBMCs | 12600000 | 1512000 | E | AEG,MD | 640010 | 80488 | 8850 | 21591 | 1356 | 14618 | 3007 | 11972 | 658 | 5 | 8369 | 10062 | 0 |
| HC_40 | M | 33.3 | CD19+ | 31800000 | 680000 | C | Mix I | 437659 | 55698 | 10944 | 30331 | 1559 | 6777 | 2617 | 1390 | 993 | 55 | 811 | 220 | 1 |
| HC_41 | F | 33.7 | PBMCs | 69000000 | 1380000 | E | AEG,MD | 1786984 | 132252 | 69315 | 31741 | 8289 | 9100 | 4499 | 2946 | 595 | 52 | 4578 | 1134 | 3 |
| HC_42 | F | 33.8 | CD19+ | 100000000 | 3360000 | C | AEG,MD | 1032835 | 195330 | 28072 | 121411 | 2521 | 23779 | 6294 | 3445 | 1021 | 139 | 4069 | 4566 | 13 |
| HC_43 | F | 34.0 | PBMCs | 3000000 | 123000 | C | AEG,MD | 1598313 | 97235 | 42852 | 38597 | 2734 | 5965 | 2844 | 1067 | 336 | 49 | 2136 | 636 | 19 |
| HC_44 | M | 35.3 | PBMCs | 15000000 | 1800000 | E | Mix I | 4584690 | 303597 | 43302 | 86951 | 11700 | 117932 | 19339 | 7425 | 3176 | 94 | 7750 | 5926 | 2 |
| HC_45 | M | 35.7 | PBMCs | 6480000 | 777600 | E | AEG,MD | 943459 | 124374 | 24273 | 36725 | 12340 | 38562 | 3062 | 4235 | 701 | 37 | 3080 | 1359 | 0 |
| HC_46 | F | 36.3 | PBMCs | 21600000 | 1300000 | E | AEG,MD | 1833493 | 187445 | 53829 | 67615 | 5050 | 20383 | 7927 | 16069 | 982 | 224 | 10156 | 5193 | 17 |
| HC_47 | M | 39.3 | PBMCs | 51000000 | 6120000 | E | Mix I | 454024 | 47055 | 5645 | 17603 | 1553 | 14413 | 3212 | 1697 | 701 | 17 | 1230 | 968 | 16 |
| HC_48 | M | 41.4 | PBMCs | 16500000 | 1980000 | E | AEG,MD | 903117 | 110233 | 32876 | 32955 | 10024 | 20751 | 2524 | 6928 | 640 | 5 | 2326 | 1204 | 0 |
| HC_49 | M | 42.6 | PBMCs | 5700000 | 684000 | E | AEG,MD | 878951 | 86541 | 26443 | 42918 | 1667 | 7504 | 2296 | 2206 | 360 | 49 | 2121 | 947 | 30 |
| HC_50 | M | 42.7 | PBMCs | 10000000 | 1200000 | E | AEG,MD | 1668134 | 69380 | 6353 | 31589 | 611 | 8976 | 3172 | 3722 | 1135 | 78 | 9926 | 3805 | 13 |
| HC_51 | F | 45.3 | PBMCs | 8400000 | 1008000 | E | AEG,MD | 1445580 | 168554 | 16932 | 37464 | 6789 | 43558 | 9925 | 27511 | 1502 | 143 | 16832 | 7879 | 19 |
| HC_52 | F | 45.4 | PBMCs | 13200000 | 1580000 | E | AEG,MD | 1694914 | 261324 | 80587 | 85933 | 15323 | 37856 | 9798 | 8977 | 4003 | 316 | 11782 | 6690 | 59 |
| HC_53 | F | 50.7 | PBMCs | 9600000 | 1152000 | E | AEG,MD | 1970119 | 116853 | 15337 | 26192 | 8018 | 32846 | 15563 | 4098 | 1241 | 47 | 8088 | 5414 | 9 |

**Supplementary table 2.**
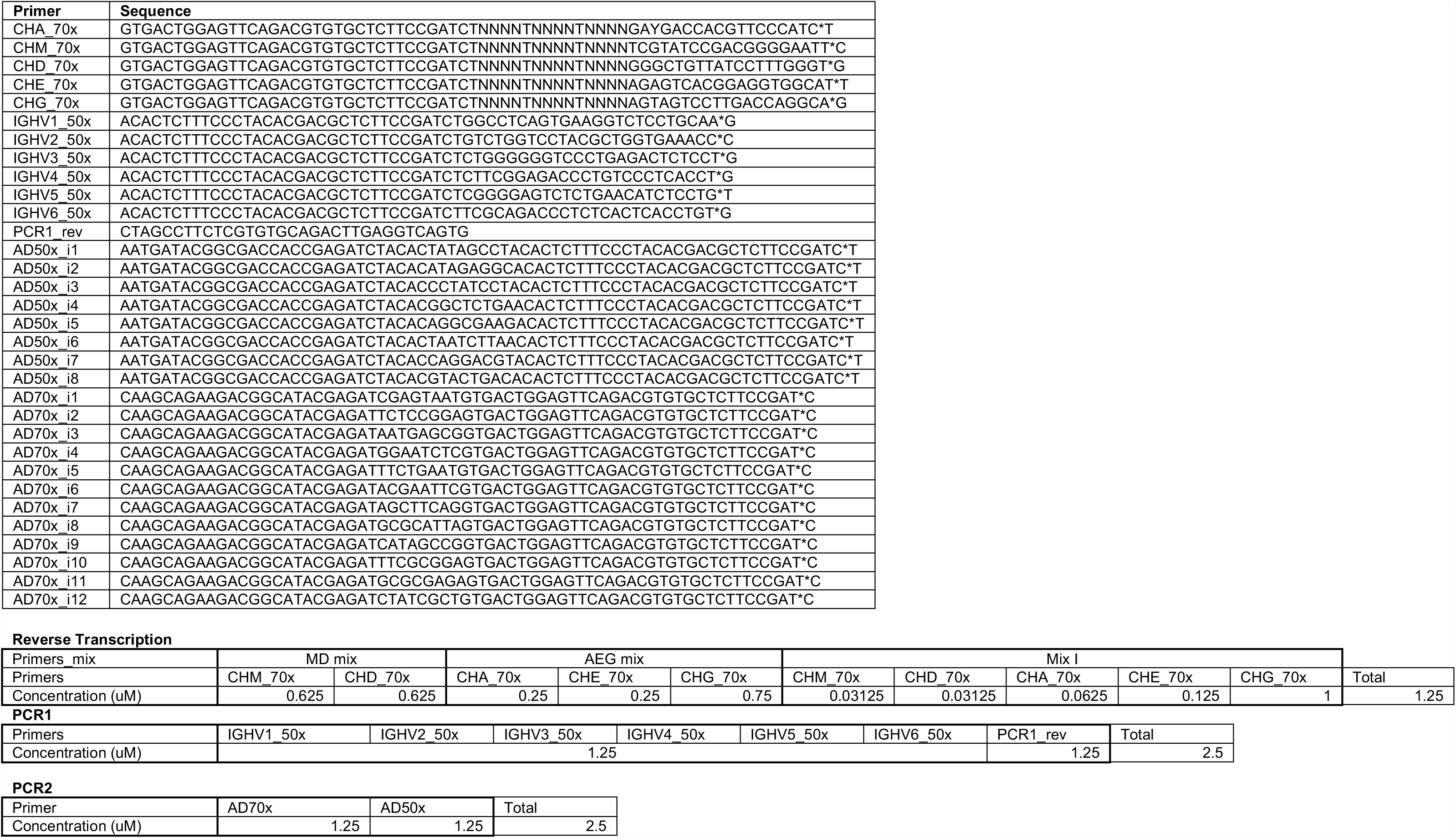

